# Scale-dependent brain age with cosmological higher-order statistics from structural magnetic resonance imaging

**DOI:** 10.1101/2025.03.24.644902

**Authors:** Aurelio Carnero Rosell, Niels Janssen, Antonella Maselli, Ernesto Pereda, Marc Huertas-Company, Francisco-Shu Kitaura

**Author notes:** Corresponding author *Email address:* (Francisco-Shu Kitaura).

## Abstract

Inferring chronological age from magnetic resonance imaging (MRI) brain data has become a valuable tool for the early detection of neurodegenerative diseases. We present a method inspired by cosmological techniques for analyzing galaxy surveys, utilizing higher-order summary statistics with multivariate two- and three-point analyses in 3D Fourier space. This method offers physiological interpretability during the inference, allowing the detection of scales where brain anatomy differs across age groups, providing insights into brain aging processes.

Similarly to the evolution of cosmic structures, the brain structure also evolves naturally but displays contrasting behaviors at different scales. On larger scales, structure loss occurs with age, possibly due to ventricular expansion, while smaller scales show increased structure, likely related to decreased cortical thickness and gray/white matter volume.

Using MRI data from the `OASIS-3` database of 869 sessions, our method predicts chronological age with a Mean Absolute Error (MAE) of 3.1 years, while providing information as a function of scale. A posterior density estimation shows that the 1-*σ* uncertainty for each individual varies between ∼ 2 and 8 years, suggesting that, beyond sample variance, complex genetic or lifestyle-related factors may influence brain aging.

We perform a twofold validation of the method. First, we apply the method to the `Cam-CAN` dataset, yielding a MAE of ∼ 5.9 years for the age range from 18 to 88 years. Second, we apply the method to thousands of simulated MRI images generated with a state-of-the-art Latent Diffusion model. This work demonstrates the utility of interdisciplinary research, bridging cosmological methods and neuroscience.

**PACS:** 87.19.lf, 87.61.Jc, 02.50.Tt, 87.19.lo

## 1. Introduction

The study of brain aging has significant implications for the early detection of neurodegenerative diseases (see, e.g., Bethlehem et al., 2022, and references therein). Understanding how the human brain evolves structurally and functionally over time can provide critical insights into age-related decline and cognitive impairment. In this work, we use “brain evolution” to mean age-related changes over the human lifespan, rather than evolutionary change across generations. Machine learning techniques, particularly those applied to large magnetic resonance imaging (MRI) datasets, have emerged as powerful tools for predicting brain age (Liem et al., 2017; Cole et al., 2017; Fisch et al., 2021; More et al., 2023; Yin et al., 2023; Meng et al., 2024). As discussed in the recent review by Kumari and Sundarrajan (2024), existing models in brain age prediction, such as ensemble methods, support vector regression (SVR), convolutional neural networks (CNN), and recurrent neural networks (RNNs), have been extensively studied and offer specific benefits depending on the task requirements.

Inspired by methods used in cosmology, we propose a novel, transversal higher-order statistical approach to characterize brain aging. In cosmology, the evolution of the cosmic web –complex structures traced by galaxies under the influence of gravity– can be described as a multidimensional distribution analyzed through higher-order statistics. Similarly, the human brain, which represents the intricate network of connections and structure in the brain, can be studied as a multidimensional distribution, with higher-order statistics revealing structural patterns across different scales. By leveraging two- and three-point statistics in Fourier space, we can capture both large- and small-scale features of the brain anatomy, offering a richer, scale-dependent understanding of how the brain ages over time.

This method does not replace machine learning techniques, but complements them. Our approach introduces a more interpretable framework by identifying the specific scales at which structural changes occur, thereby enhancing the robustness of machine learning predictions.

Moreover, our method provides two key advantages—higher-order statistics (Hindriks et al., 2024) and flexibility to choose across scales—that can be directly compared with traditional neuroscience approaches (Bethlehem et al., 2022; Oschwald et al., 2019; Yang et al., 2024). While conventional methods, such as assessing hippocampal volume and ventricular size or analyzing saliency maps (Yin et al., 2023), have advanced our understanding of gross structural changes in the aging brain, they often miss subtle changes governed by the nonlinear dynamics of structural relationships across scales. These intricate dynamics can be captured more specifically using multidimensional higher-order statistical Fourier analysis, as demonstrated in this work. By integrating these techniques, we can accurately relate individual brain structure to biological age and potentially gain deeper insight into the physiological processes underlying brain aging, improving early diagnosis of neurodegenerative diseases.

## 2. Theoretical and mathematical background

Statistical analysis of fields in cosmology often requires understanding their underlying probability distribution functions (PDFs) and their deviations from Gaussianity. These deviations are quantified using moments and cumulants, which describe statistical properties such as variance, skewness, and higher-order interactions. In this section, we introduce the connection between correlation functions, moments, and cumulants, starting from a general non-Gaussian PDF (see, e.g., Bernardeau et al., 2002; Kitaura, 2012, and references therein). For a modern reference to multivariate statistical methods see Pemantle et al. (2024).

We model observables as realizations of a non-Gaussian random field (the MRI scan) defined over a three-dimensional domain. Two standard assumptions simplify the statistics:

- **Statistical homogeneity** (translation invariance): ensemble averages do not depend on absolute position.
- **Statistical isotropy** (rotation invariance): in the absence of preferred directions, correlation functions depend only on separations.

These assumptions call for an extension of this work including anisotropic clustering analysis, which we leave for future studies (see also the discussion at the end of Appendix F). For a purely Gaussian field, the two-point statistics—i.e., the correlation function *ξ*(*r*) (a function of distance *r*) or its Fourier counterpart, the power spectrum *P* (*k*) (a function of wavenumber. inverse of distance, *k*)—completely characterize the signal; all connected moments of order *n >* 2 vanish. We detail these quantities below. By contrast, cosmological structure formation, nonlinear evolution, and astrophysical processes generate departures from Gaussianity, motivating the use of cumulants and higher-order spectra, such as bispectra. An analogous situation holds for the healthy human brain: its organized, anisotropic, and modular architecture is far from featureless noise, so higher-order statistics are likewise necessary to capture intrinsically multi-point dependencies.

### 2.1. Moments and cumulants of a non-Gaussian PDF

Consider a multivariate probability distribution function *P* (***ν***) describing the statistical distribution of a field *I*(**x**), where **x** is a three-dimensional coordinate (e.g., the voxel center). The variable *ν* is typically a function of *I*(**x**), such as *ν* = *I* or *ν* = *I/* ⟨*I*⟩ − 1, the latter being a zero-centered quantity. The *n*-th order *moments* are defined as:

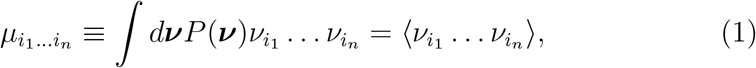

The moment generating function (MGF) is defined as:

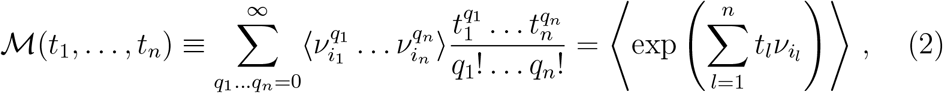

and hence, ℳ_*ν*+*η*_(*t*) = ℳ_*ν*_(*t*)ℳ_*η*_(*t*) (for independent random variables *ν* and *η*). Subsequent derivatives of ℳ(*t*_1_, …, *t*_*n*_) at the origin **t** = 0 yield the moments:

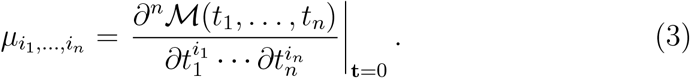

Moments capture the overall shape of a distribution but do not distinguish between independent contributions and genuinely correlated structures. To isolate connected correlations, we define cumulants, which are derived from the cumulant generating function (CGF):

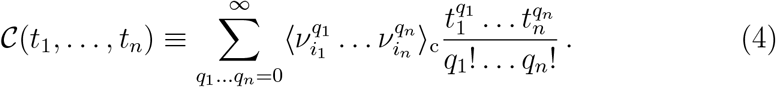

The cumulants 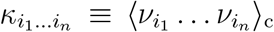, where the subscript *c* stands for the connected part, are obtained by taking derivatives of 𝒞(*t*_1_, …, *t*_*n*_) with respect to the parameters *t*_*i*_. Specifically, the *n*-th order cumulant is given by:

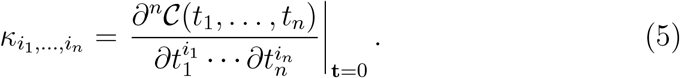

Since cumulants isolate genuine correlations without mixing information from independent and dependent sources, they must satisfy the additive property for independent random variables: *κ*_*n*_(*ν* + *η*) = *κ*_*n*_(*ν*) + *κ*_*n*_(*η*).

This property is already exhibited by measures such as the mean and variance for independent variables. Higher-order statistical dependencies should also obey a similar rule if they are to serve as meaningful descriptors of intrinsic structure. Since cumulants are obtained from the derivatives of the CGF, the CGF itself must also satisfy this additivity property: 𝒞_*ν*+*η*_(*t*) = 𝒞_*ν*_(*t*) + 𝒞_*η*_(*t*)

This property suggests that the relationship between the CGF and the MGF follows a logarithmic transformation:

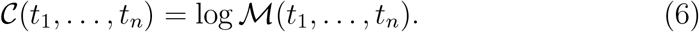

This transformation ensures that multiplicative relationships between MGFs translate into additive relationships between CGFs.

By combining equations 2, 4 and 6 we obtain:

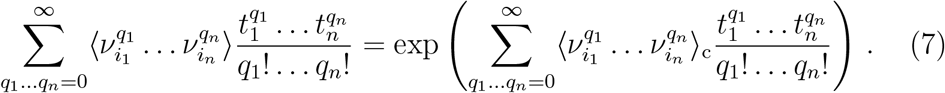

Hence, moments can be expressed as a sum of cumulants, which can be realized by looking at all combinations of connections between points in a diagram, as shown in Figure 1.

**Figure 1.**
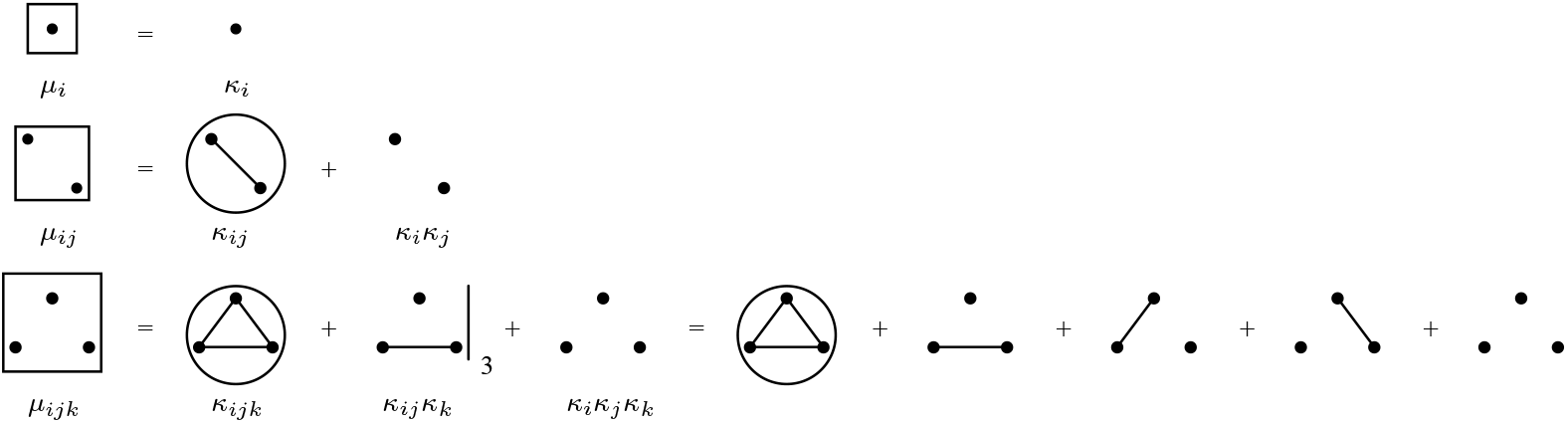
Diagrammatic representation of moments up to third-order correlation statistics (where we choose for simplicity the notation: *i* ≡ *i*_1_, *j* ≡ *i*_2_, *k* ≡ *i*_3_ and each index runs over all voxels). The rectangular boxes denote ensemble averages, corresponding to statistical moments. In the first row we have the mean (in the univariate case: 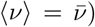), in the second row we have the relationship for the variance (in the univariate case: 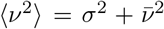). Dots represent individual data points, with connected dots indicating cumulants (connected moments). The numbers next to certain configurations indicate the count of equivalent terms. Circles highlight the terms that are nonzero, as they lack fully independent components when assuming a centered variable with zero mean. The three equivalent terms of the third-order moment are explicitly shown. A ‘cumulant object’ is defined as a set of connected points within a moment term, with two cumulant objects considered equivalent if they contain the same number of points.

The second-order cumulant corresponds to the two-point correlation function:

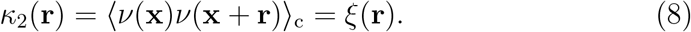

Similarly, the third-order cumulant corresponds to the connected part of the three-point correlation function:

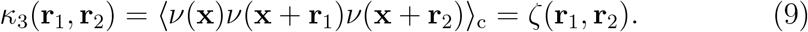

Higher-order cumulants describe more complex interactions and deviations from Gaussianity. Expanding this function systematically isolates the connected statistical contributions, removing contributions from independent fluctuations.

### 2.2. Power spectrum and the two-point correlation function

The power spectrum *P* (**k**) characterizes the two-point statistical properties of a field *ν*(**x**) with **k** being the Fourier three-dimensional k-vector. Given its Fourier transform,

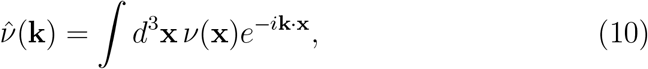

the power spectrum is defined as

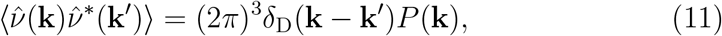

where *ν*^∗^(**k**) is the conjugate of *ν*(**k**), and *δ*_D_ the Dirac delta-function. In numerical computations, the power spectrum is estimated as:

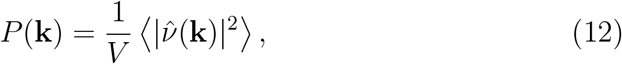

where *V* is the survey volume, and the averaging is performed over bins in Fourier space.

The power spectrum is directly related to the two-point correlation function *ξ*(**r**), which measures the spatial correlations of a field in real space:

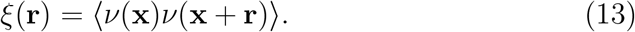

The connection between the power spectrum and the two-point correlation function is given by the Fourier transform pair (according to the so-called Wiener-Khinchin theorem: Wiener, 1930; Khintchine, 1934):

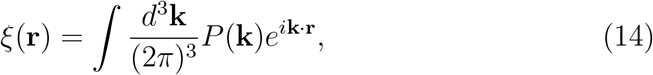

which illustrates how the power spectrum provides information about the clustering properties of the field in Fourier space.

### 2.3. Bispectrum and the three-point correlation function

The bispectrum measures the three-point correlations of the Fourier-transformed field and is defined as

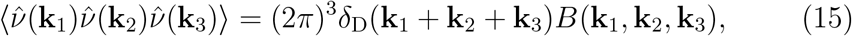

where the Dirac delta function enforces the triangle closure condition, **k**_1_ + **k**_2_ + **k**_3_ = 0.

The bispectrum is related to the three-point correlation function *ζ*(**r**_1_, **r**_2_), which quantifies the probability of finding triplets of points at separations **r**_1_ and **r**_2_:

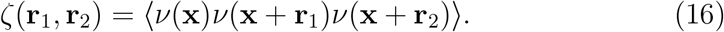

The Fourier transform relationship between the bispectrum and the three-point correlation function is given by:

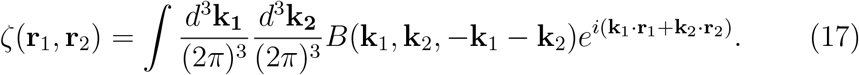

This highlights how the bispectrum captures non-Gaussian features that are not fully described by the power spectrum alone.

### 2.4. Hierarchical model for higher-order correlation functions

Hierarchical models assume that structures exhibit self-similarity, allowing higher-order correlation functions to be constructed from products of the two-point correlation function (Fry and Peebles, 1978; Fry, 1984b,a; Bardeen et al., 1986; Balian and Schaeffer, 1989):

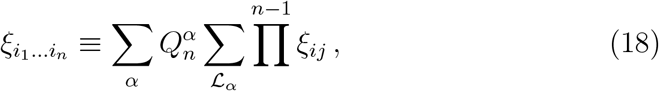

such that the full set of points *i*_1_ … *i*_*n*_ is connected by links of *ξ*_*ij*_. These links form a tree structure, where *α* represents different tree topologies for each order *n*. The sum over ℒ_*α*_ accounts for all possible labelings of a given tree. The hierarchical coefficients 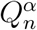 encode the remaining degrees of freedom.

For the three-point correlation function, the hierarchical model leads to:

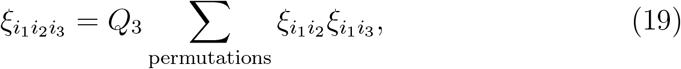

where *Q*_3_ is the single hierarchical coefficient at third order.

In Fourier space, the normalized (*Q*_3_ = 1) hierarchical bispectrum is defined as:

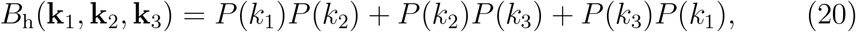

which leads to the definition of the reduced bispectrum, highlighting non-Gaussian deviations:

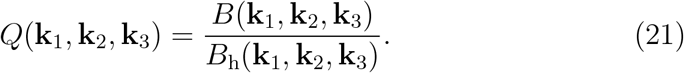

This formulation accounts for expected Gaussian-like hierarchical structures and isolates deviations from Gaussianity.

## 3. Method

Within the framework of information theory, the set of voxels in a three-dimensional image can be interpreted as a probability distribution function, shaped by an underlying physical and observational process.

An image, such as an MRI image, is always defined over a finite, positive range. The intensity distribution from such images, modeled by a compactly supported PDF, is constrained to a bounded domain. Despite this finite support, the complexity of the underlying intensity distribution can be arbitrarily high, reflecting intricate anatomical or (physio-)pathological structures. However, the advantage of distributions with finite support is that their moment sequences are often better behaved, and the information encoded in these moments is more constrained. In many cases, the moments uniquely determine the distribution—a property known as moment determinacy—without requiring additional regularisation or assumptions that are necessary for distributions with infinite or unbounded support (Stoyanov et al., 2020). This makes finite-support distributions particularly suitable for moment-based analysis (Hausdorff, 1921a,b; Shohat and Tamarkin, 1943).

Since we are dealing with three-dimensional data we have to consider correlation functions of multivariate PDFs, for which each voxel of the MRI image represents one statistical dimension.

We compute spatial correlations of the scalar field *I*(**x**) obtained from the preprocessed structural MRI (bias-field corrected, skull-stripped, and intensity-normalized within the brain mask). Importantly, we do *not* use tissue-probability maps or modulated volumes; the analysis is performed directly on *I*(**x**).

Analogously to the two-point correlation function of galaxies, which is a function of distance and describes the excess probability of finding two galaxies separated by this distance (Peebles, 1980), we consider the correlation of voxels based on their intensities. This is computed in Fourier-space, yielding the so-called power spectrum (see section 2 and Vazza and Feletti, 2020, for a power spectrum study of dead brain structures). Distributions with identical two-point statistics can have very different three-dimensional patterns (Kitaura et al., 2015). Thus, we need to resort to higher-order statistics.

The three-point correlation function has a long history in cosmological astronomy. It was already computed in configuration space in the 70s from the three-dimensional galaxy distribution (Groth and Peebles, 1977). Later, it was extended to Fourier-space, the so-called bispectrum (Baumgart and Fry, 1991). The three-point statistics can be exploited to probe models of structure formation (Frieman and Gaztanaga, 1994), the bias from dark matter tracers (Matarrese et al., 1997), or neutrino masses (Hahn and Villaescusa-Navarro, 2021). It has the powerful property of capturing deviations from a Gaussian distribution, and can thus be extremely sensitive to the complex multidimensional patterns beyond the two-point statistics. Controlling both two- and three-point statistics of complex structures provides a powerful approach for accurate modeling. For example, the distribution of galaxies and their dynamics are successfully reproduced by constraining model parameters using two- and three-point statistics, as demonstrated in previous work where the same software used here was applied to galaxy survey data (Kitaura et al., 2016).

In this study, as part of the `Cosmic Brain` project^1^, we rely on the combined power spectrum and bispectrum analysis (see section 2, Figure 2 and corresponding explanation in Appendix B). In Fourier-space, signals are decomposed into different frequency components, to different spatial scales. This allows for a more detailed and clear understanding of how structural changes manifest at different scales. For brain anatomy, this is crucial because aging or neurodegenerative processes impact large-scale (global) and small-scale (local) brain structures differently, as we will show below. A detailed description of the method presented in this work can be found in Appendix D.

**Figure 2.**
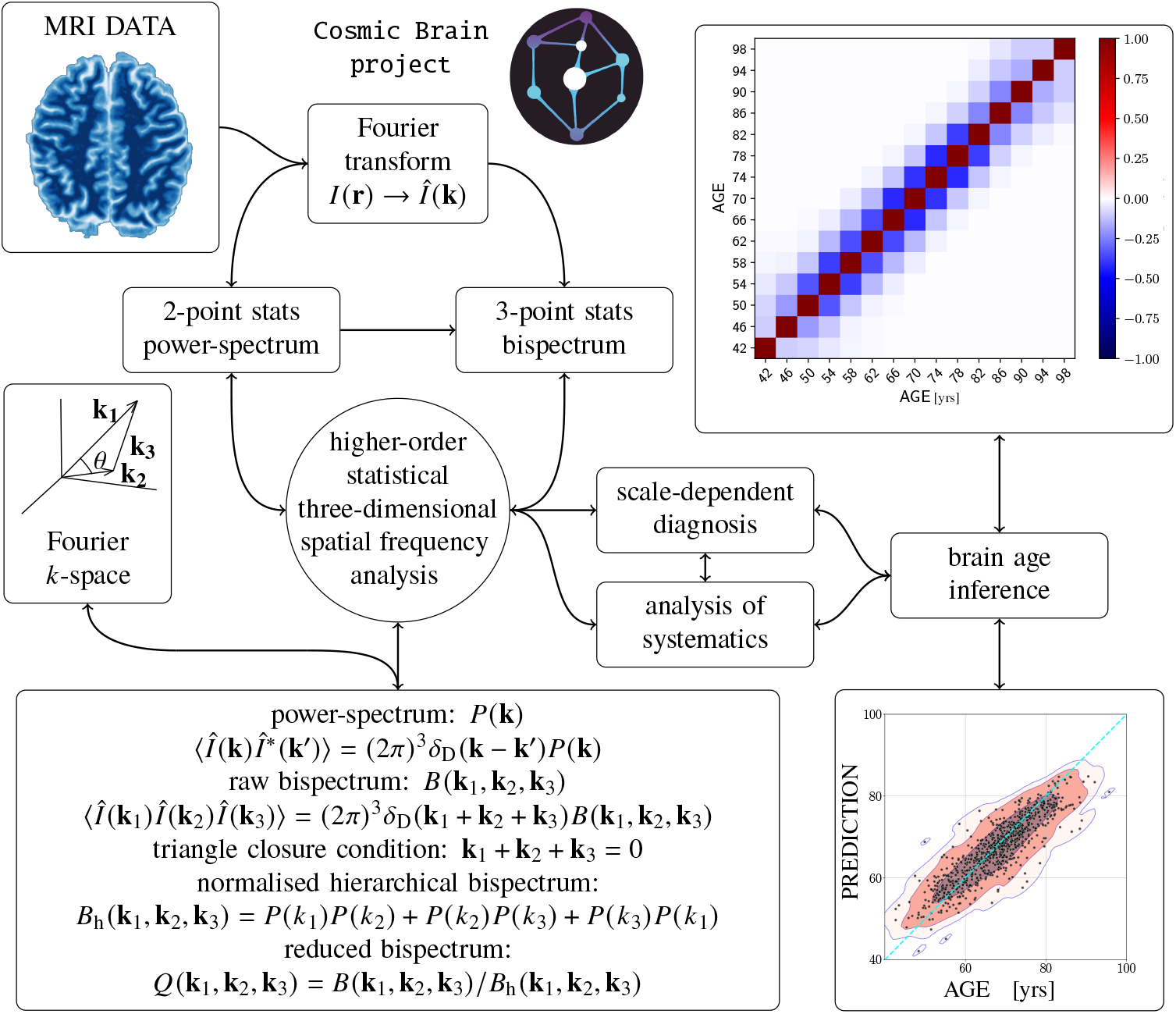
Summary of the higher-order statistical three-dimensional spatial frequency analysis method for brain age inference (as part of the `Cosmic Brain project`), showing the process from the input MRI data (top-left) to the outcome (right): normalized covariance matrix (top-right) and regression between the predicted brain age and the actual chronological age (bottom-right). The MRI data can be represented as an intensity array *I*(**r**) in a cubical mesh with voxel coordinates **r** = {*r*_*x*_, *r*_*y*_, *r*_*z*_}. The Fourier transform of *I*(**r**) is represented by 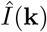 as a function of the *k*-vector. The power spectrum *P* (**k**) represents a measure of the two-point clustering statistical (squared) amplitude as a function of spatial frequency distance, while the bispectrum *B*(**k**_1_, **k**_2_, **k**_3_) represents the analogous quantity for the three-point statistics, as a function of triangle configurations. 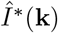 is the conjugate of 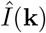, and *δ*_D_ is the Dirac delta function.

### The OASIS-3 dataset

`OASIS-3` MRI data were collected through the Knight Alzheimer’s Disease Research Imaging Program at Washington University in St. Louis, MO, USA (LaMontagne et al., 2019), using a Siemens Vision 1.5T scanner or, for most participants, two versions of a Siemens TIM Trio 3T scanner (Siemens Medical Solutions USA, Inc.). Participants were scanned in the supine position, with head motion minimized using foam pads, and a vitamin E capsule was placed over the left temple for lateralization in some cases. A 16-channel head coil was used for all scans. For the present study, we analyzed only T1-weighted structural images, acquired with a 3D MPRAGE sequence (TI/TR/TE = 1000/2400/3.08 ms, flip angle = 8°, 1 mm isotropic voxels). Although the `OASIS-3` dataset includes additional imaging modalities, such as FLAIR, DTI, and ASL, these were not considered in this analysis.

Our dataset consists of 869 MRI sessions (see Appendix A for details on sample preparation and selection) from 378 unique “healthy” subjects in the `OASIS-3` database. From now on, we will refer to each MRI session as a subject, for simplicity, using the age recorded during the session as the subject’s age on the day of data acquisition.

The `OASIS-3` sample includes subjects aged 42 to 98 years, though the distribution is notably sparse below 50 and above 85.

## 4. Results

We calculate the power spectra, *P* (*k*), and bispectra for 975 triangular configurations, thus measuring the two- and three-point statistics in Fourier-space. In particular, we run over triangle configurations given the length of two sides as a function of the subtended angle, *θ*. Considering the size of human brains and the resolution of the MRI data we consider the spatial frequency range from *k* = 0.1 mm^−1^ to *k* = 2 mm^−1^ in Fourier-space, equivalent to *r* ≈ 3 mm and *r* ≈ 63 mm in configuration space.

We include two types of bispectra in the analysis: raw, *B*(*θ*), and reduced, *Q*(*θ*) (see section 2). Since the reduced bispectrum is computed as the raw bispectrum divided by the hierarchical bispectrum model, which is constructed from the sum of permutations of power spectrum products, a combined analysis of raw and reduced bispectra effectively integrates information from both two-point and three-point statistics.

### 4.1. Brain age dependence on the higher-order statistics

We find that the power spectrum of intensities shows a nearly linear, moderate correlation of amplitude with age (see lower-left panel in Figure 3), suggesting limited potential as a standalone biomarker for brain age prediction. In contrast, the bispectra exhibit clear, complex, and strong correlations with age (see middle and right panels in the same figure).

**Figure 3.**
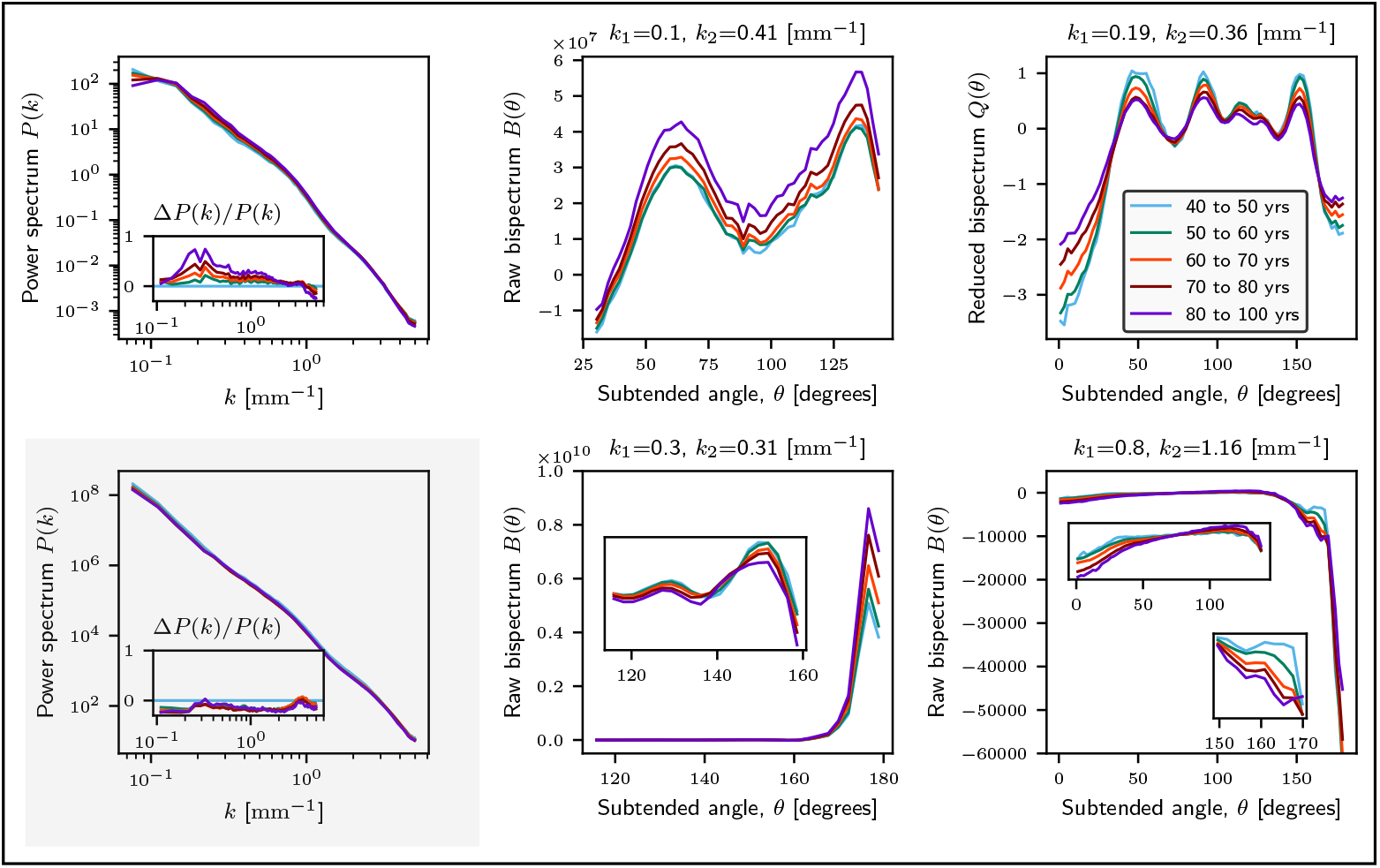
Median power spectra per age group: upper left-panel: setting average intensity outside the scanned region, lower left shaded-panel: setting zero intensity outside the scanned region. Middle and right panels: bispectra per age group for the individual configurations that enter the biomarker definition. Here we consider the raw and reduced bispectra for fix |**k**_1_| and |**k**_2_| as a function of the subtended angle *θ* between both *k*-vectors: *B*(*θ*|*k*_1_, *k*_2_) and *Q*(*θ*|*k*_1_, *k*_2_).

To enhance the power spectrum signal, we introduce a mask treatment very common in cosmology, which consists of working with fluctuations along the mean intensity and assuming average intensity outside the region. The resulting power spectra accentuate the age evolution (see upper-left panel in Figure 3).

The ordering of the curves by age groups highlights the potential of this technique as a tool for diagnosing neurodegenerative diseases. We find that reduced bispectra are more sensitive on larger scales, while raw bispectra are more sensitive to small scales. This behaviour is quantified through Kendall’s rank correlation coefficient *τ* (Kendall, 1938, 1945) shown in Appendix C, which is better suited than Pearson’s correlation, as it does not assume a linear relationship, but evaluates how well the ranks of one variable correspond to the ranks of another variable.

Finally, since the complexity in the three-dimensional structure can be characterized by the deviation from a flat bispectrum, we propose the use of the variance as a way to identify outliers. In fact, it helped us to identify outliers that were subsequently removed from the analysis (see Appendix A.2 for details on the selection criteria). These outliers may result from MRI measurement artifacts, early signs of dementia, or general anatomical anomalies, which are referred to as systematics in the context of our study. In future studies, with more data available, we plan to investigate these outliers in more depth to determine whether they are due to systematics or represent healthy individuals with unique brain evolution linked to particular lifestyle factors.

### 4.2. Statistical analysis

First we evaluate the pool of 975 triangle configurations. To estimate the performance of each configuration as a potential biomarker, we employed a Random Forest classifier (see Breiman, 2001) using a leave-one-out cross-validation (LOO-CV) technique (Hastie et al., 2001), ensuring that each subject’s estimate was independent of the training sample.

To assess the classification performance, we calculate the MAE and the geometric mean score (G-Mean, defined as the square root of the product of class-wise sensitivity, see Appendix C for details), both as overall averages and as functions of age.

A practical biomarker should not rely on ∼ thousands of computationally expensive Fast Fourier Transform (FFT) calculations. Additionally, many of these configurations likely contain redundant information or exhibit weaker correlations with age compared to others. Therefore, it is essential to reduce the number of informative configurations to a small subset that can be computed within a few minutes per patient for potential clinical applications, without compromising diagnostic accuracy.

To this end, we identified the four top singular configurations (see Figure 3 and Table 1) and we integrated them into an age-regression neural network (see Appendix D for further explanations on the algorithm).

**Table 1:** Summary of singular configurations making the OASIS-3 biomarker.

| | $k_1$ | $k_2$ | $r_1$ | $r_2$ | $\theta$ | CPU time | Name |
| --- | --- | --- | --- | --- | --- | --- | --- |
|  | [mm<sup>-1</sup>] |  | [mm] |  | [rad] | [s] |  |
| *B* | 0.10 | 0.41 | 62.83 | 15.32 | 0.5-2.5 | 2 | B010041 |
| *Q* | 0.19 | 0.36 | 33.07 | 17.45 | 0.-3.15 | 4 | Q019036 |
| *B* | 0.30 | 0.31 | 20.94 | 20.27 | 2.-3.15 | 5 | B030031 |
| *B* | 0.80 | 1.16 | 7.85 | 5.43 | 0.-3.15 | 660 | B080116 |

Table 1 also reports run times for each bispectrum configuration using a single CPU. Large-scale (low-*k*) triangles are computed efficiently, whereas small-scale (high-*k*) configurations are substantially more expensive. In practice, this makes low-*k* workloads viable for on-device use (e.g., a smartphone app), while high-*k* evaluations remain impractical on current mobile hardware. We anticipate substantial speedups from our new GPU implementation in the cosmological setting (Rosselló et al., 2025), which should narrow this gap.

The noisy patterns observed in some of the raw bispectrum configurations, suggest that there is potential for improvement with the availability of more data. It is important to note, however, that lifestyle factors increasingly influence biological brain age as individuals grow older. Nonetheless, we plan to explore the completeness of the information in the summary statistics by extending this study to smaller-scale bispectrum configurations and the trispectrum (four-point statistics), while also conducting brute-force analyses on using the full cubical MRI data volumes (see, e.g., methods such as García-Farieta et al., 2024).

We apply the analysis to the `OASIS-3` sample of 869 sessions. The bottom-right panel of Figure 2 and both panels in Figure 4 presents the scatter plot comparing chronological age with the predicted age, as obtained using the age-regression neural network developed in this study. In addition, the top-right panel of Figure 2 shows the normalized covariance matrix (see, e.g., Kitaura et al., 2016) for age groups in four-year intervals.

**Figure 4.**
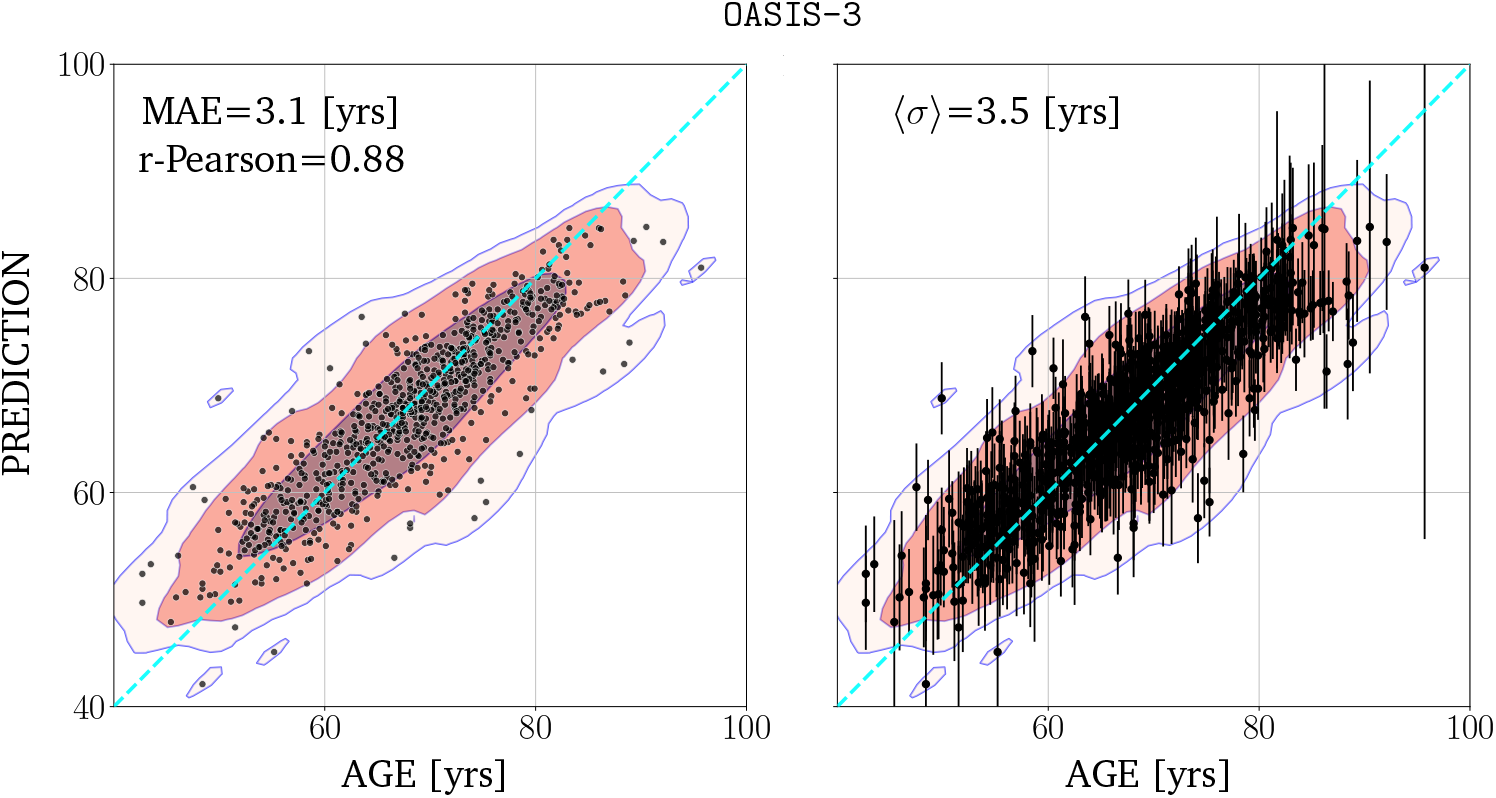
Regression based on the combined two- and three-point statistics according to Table 1 applied on the `OASIS-3` dataset using deep ensemble samples indicating (on the left:) the MAE and *r*-Pearson correlation coefficient; and (on the right:) the standard deviation per subject averaging over 1000 samples.

We estimate posterior densities for age using 1000 realizations, and report the posterior mean and posterior standard deviation as the point estimate and associated uncertainty, respectively. In future work, we aim to explore the calibration and robustness of the estimated posteriors in greater detail with larger datasets, to obtain more precise uncertainty quantification for each patient. The covariance matrix exhibits a strong diagonal structure with rapidly diminishing off-diagonal elements. This pattern is indicative of low redundancy among features and a well–conditioned inverse covariance, which would be consistent with tight age constraints under a Gaussian–likelihood approximation (Cramér, 1946; Rao, 1992). In the present work, we report this covariance only as an exploratory diagnostic: it is neither used as input to the regressor nor propagated into our error bars. A full covariance–based uncertainty analysis is deferred to future work.

The overall average MAE across 1000 deep ensemble samples for the `OASIS-3` dataset is 3.1 years. The average standard deviation yields 3.5 years, closely aligning with the MAE and confirming the robustness of the analysis.

We also tested the inclusion of the power spectrum (upper left panel in Figure 3) in combination with the raw bispectra, replacing the reduced bispectrum `Q019036`. This configuration yielded a MAE of 3.3 years. These results suggest that, for this dataset, the reduced bispectrum may encode more information than the power spectrum in combination with a reduced number of raw bispectra.

### 4.3. Validation of the method

To validate our approach, we first applied the methodology to simulated data (see Appendix E). We then confirmed that the age inference results were consistent with those reported in the original study by Pinaya et al. (2022). Notably, our method outperformed the reference model—a 3D convolutional neural network—, achieving a higher Pearson correlation coefficient (*r* = 0.77 vs. *r* = 0.69; see Figure 3 in Pinaya et al. 2022 for comparison). The resulting age regression is illustrated in Figure E.8. Alternative methods to generate simulated data can be taken into account in future studies (see, e.g., Wen et al., 2022; Yang et al., 2022; Bhattarai et al., 2024).

Second, we applied the methodology to the `Cam-CAN CC700` dataset (Taylor et al., 2017; Shafto et al., 2014, see Appendix F for details). Even though the characteristics of this dataset is significantly different from `OASIS-3` we obtain qualitatively similar results (MAE for the age range: 18-88: ∼ 5.9, see right panel in Figure F.12 for a comparison), considering the sparser dataset across a wider age range. In particular, we find that some of the bispectrum configurations obtained from `OASIS-3` are still valid in `Cam-CAN`, even though they sample different age ranges with different densities and populations. For this dataset, the method preferred the use of the power spectrum, instead of the reduced bispectrum `Q019036`, as well as another configuration tracing smaller scales, B066107, instead of the large scale configuration B010041.

## 5. Neuroanatomical and physiological interpretation

We present a correlation study between the bispectrum configurations and the neuroanatomical variables of the brain for two configurations, `Q019036` and `B080116`. In particular, we focus on Kendall’s rank-correlation. Nonetheless, we have performed also a p-value mutual information calculation which confirms Kendall’s analysis results (see Figure 5).

**Figure 5.**
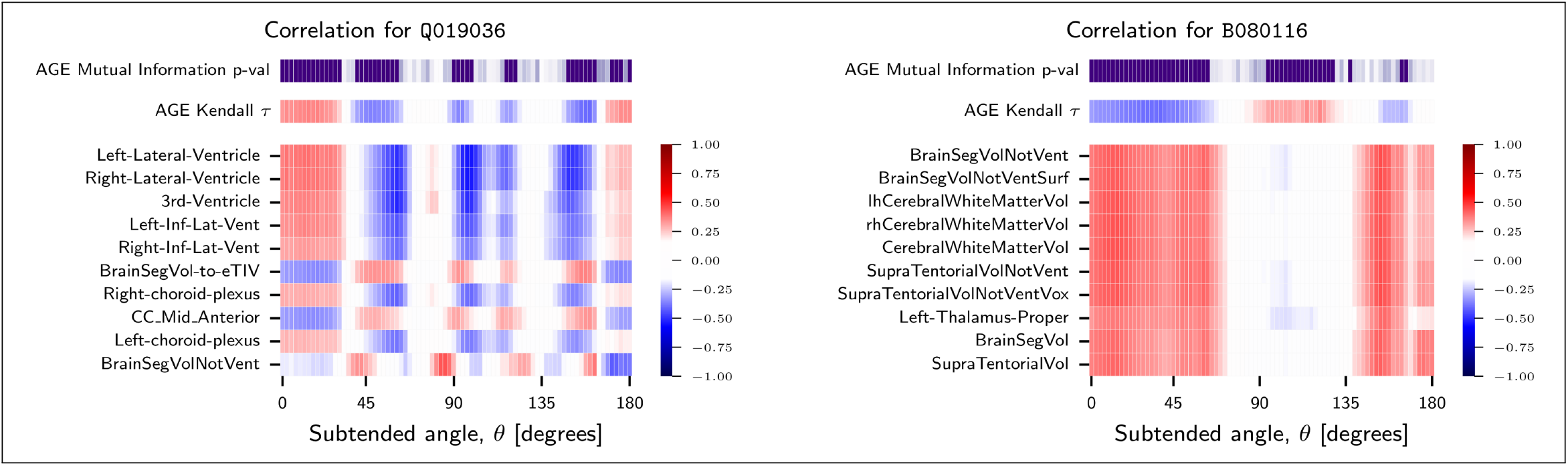
At the top of both panels, p-values from mutual information analysis for bispectra across subtended angles with age (purple color indicates strong dependence: *p <* 0.001). Below that, Kendall’s rank correlation to the bispectra for two configurations are shown: with age (top) and with the ten most significant anatomical variables from Freesurfer. In particular, at the scales of Q019036, the effect of ventricle volumes is highlighted, while at smaller scales in B080116, the bispectrum is more sensitive to white matter content. Red indicates positive correlation, while blue represents negative correlation or anti-correlation.

### 5.1. Scale-dependent analysis

Since the bispectrum configuration is determined by two wavenumbers, *k*_1_ and *k*_2_, which define the sides of a triangle in Fourier space, it captures interactions across different spatial scales. Considering that the Fourier wavenumbers are inversely proportional to the scale:

- **Small** *k*_1_, *k*_2_ **values**: Correspond to large-scale structures, capturing global brain morphology changes, such as cortical shrinkage and ventricular expansion.
- **Large** *k*_1_, *k*_2_ **values**: Reflect fine-scale structures, highlighting local variations in cortical thickness, white matter integrity, and microstructural degradation.
- **Intermediate configurations**: Represent mixed-scale interactions, where structural correlations exist between global and local features.

#### Bispectrum angles

The angular relationships between *k*_1_ and *k*_2_ further refine the interpretation, with small angles representing interactions between similar-scale structures and intermediate to large angles reflecting multi-scale dependencies. The angle *θ* between *k*_1_ and *k*_2_ defines how different scales interact within a bispectrum triangle:

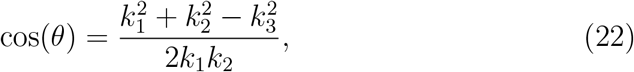

according to the law of cosines, where *k*_3_ is the third side of the triangle, constrained by the triangle inequality. The angular dependence plays a crucial role in the interpretation of bispectrum correlations:

- **Small angles (***θ <* 30^*°*^): These correspond to nearly collinear wavevectors (*k*_1_ ≈ *k*_2_), meaning the bispectrum predominantly captures self-similar interactions at the same scale.
- **Intermediate angles (**30^*°*^ *< θ <* 170^*°*^): These angles represent configurations where *k*_1_ and *k*_2_ are significantly different, leading to cross-scale interactions that mix structures of smaller and larger scales.
- **Very large angles (***θ >* 170^*°*^): These correspond to nearly antiparallel wavevectors, meaning *k*_1_ and *k*_2_ point in opposite directions but maintain similar magnitudes. This configuration behaves similarly to small angles because it still emphasizes self-similar interactions, but mirrored across larger spatial extents.

This symmetry in behavior between small and very large angles arises from the geometry of Fourier interactions: when wavevectors are nearly collinear or anti-parallel, the dominant structural patterns remain confined to closer-scale interactions, whereas intermediate angles introduce scale-mixing effects that modify the correlation patterns. In particular, the bispectrum in the squeezed limit, i.e., for *k*_3_ ≪ *k*_1_, *k*_2_, can be expressed in a more compact form and is widely studied in cosmology (see, e.g., Creminelli et al., 2011; Pajer et al., 2013).

It is important to note that, in general, we are considering configurations where *k*_1_ and *k*_2_ are significantly different, except in the case of `B030031`. When *k*_1_ ≃ *k*_2_, the bispectrum in the squeezed limit admits a simpler expression that depends solely on a function of the power spectrum.

Additionally, we distinguish between reduced and raw bispectra. Thus, by analyzing the correlation between bispectrum configurations and anatomical features, we are studying how different spatial interactions in the brain relate to specific neuroanatomical structures.

### 5.2. Bispectrum, Kendall’s rank correlation and mutual information analysis

Our bispectrum analysis, combined with Kendall’s rank correlation, and mutual information calculations, provides further insights into how structural changes in the brain manifest across different scales and configurations. We identified key differences in correlations based on bispectrum angles and wavenumber configurations.

#### For Small angles (*θ* < 30^*°*^) and very large angles (*θ* > 170^*°*^)

- **Large-scale interactions: reduced bispectrum configuration (***k*_1_ = 0.19, *k*_2_ = 0.36):
  - Strong positive correlations with ventricular volumes (left and right lateral ventricles, third ventricle, inferior lateral ventricles), suggesting that large-scale morphological changes such as ventricular expansion are major drivers of non-Gaussian features in brain aging.
  - Negative correlations with total brain volume measures (e.g., Brain-SegVol-to-ETIV, corpus callosum mid-anterior), reinforcing the interpretation that structural degradation and atrophy reduce overall brain complexity.
- **Small-scale interactions: raw bispectrum configuration (***k*_1_ = 0.80, *k*_2_ = 1.16):
  - High positive correlation with white matter and supratentorial volume measures, implying that fine-scale cortical and white matter integrity strongly contribute to bispectrum variability at these scales.
  As age increases, the raw bispectrum captures an increase in structural complexity at small scales, possibly due to cortical folding compensation mechanisms as white and gray matter decline.

### 5.3. Interpreting the brain structures in relation to bispectrum features

The brain regions analyzed can be broadly classified into ventricular structures, cortical structures, and volume-based measures, each having distinct physiological significance:

#### 5.3.1. Ventricular System (CSF Spaces)

- **Left-Lateral Ventricle, Right-Lateral Ventricle, 3rd Ventricle, Left-Inf-Lat-Ventricle, Right-Inf-Lat-Ventricle**
  - These structures contain cerebrospinal fluid (CSF) and expand with brain atrophy (e.g., in neurodegenerative diseases).
  - Strong bispectrum correlations with these ventricles may indicate that specific scales of non-Gaussianity are linked to ventricular enlargement, which is a biomarker for neurodegeneration (e.g., Alzheimer’s, hydrocephalus).
  - For small angles (*k*_1_ = 0.19, *k*_2_ = 0.36) in the reduced bispectrum, Kendall’s rank correlation is high for the Left-Lateral Ventricle, Right-Lateral Ventricle, 3rd Ventricle, and Left-Inf-Lat-Ventricle, showing a strong positive relationship. Hence, ventricular structures are well captured by large scale (low *k*-mode) bispectrum configurations.
  - The correlation is slightly lower but follows the same trend for the Right-Inf-Lat-Ventricle, right-/left-choroid-plexus.
  - For intermediate angles, the correlation reverses, approaching -1 for these ventricular structures. This strong anticorrelation suggests that bispectrum features at these scales are sensitive to ventricular expansion but in an inverse manner: larger ventricles are associated with lower bispectrum amplitudes in these configurations. This pattern may indicate a shift in the dominant spatial patterns of non-Gaussianity, potentially reflecting anatomical remodeling or differential sensitivity to tissue-CSF contrast at intermediate scales.

#### 5.3.2 Brain Volume and Cortical Measures

- **BrainSegVol-to-ETIV (Brain Segmentation Volume to Estimated Total Intracranial Volume)**
  - This ratio is a normalized brain volume measurement, adjusting for head size.
  - Correlations with bispectrum suggest that certain spatial scales of non-Gaussianity are predictive of brain volume loss.
  For small angles in the reduced bispectrum (*k*_1_ = 0.19, *k*_2_ = 0.36), Kendall’s rank correlation is negative, suggesting that increased bispectrum strength is associated with a reduction in brain volume. This implies that large-scale bispectrum features are primarily capturing structural atrophy patterns, where a decrease in brain volume corresponds to an amplification of non-Gaussianities at these scales.
  - At intermediate angles, the correlation flips to approximately 1, indicating that bispectrum features at these scales track brain volume in an opposite manner. This suggests that at intermediate Fourier modes, non-Gaussian structure may be reflecting localized cortical thinning, tissue heterogeneity, or compensatory spatial reorganization rather than global atrophy, leading to an inverse dependence on bispectrum strength compared to larger scales.

#### 5.3.3. White Matter and Supratentorial Volumes

- For the raw bispectrum configuration with *k*_1_ = 0.80 and *k*_2_ = 1.16:
  - For both small and large angles, Kendall’s rank correlation is high for:
    * **BrainSegVolNotVent:** Measures total brain volume excluding ventricles.
    * **BrainSegVolNotVentSurf:** Similar to BrainSegVolNotVent but calculated using surface-based methods.
    * **Left and Right Cerebral White Matter Volumes (lh/ rhCerebralWhiteMatterVol):** Quantifies the volume of white matter in each cerebral hemisphere.
    * **CerebralWhiteMatterVol:** Total cerebral white matter volume, combining left and right hemispheres.
    * **SupratentorialVolNotVent/Vox:** Measures the volume of the brain above the tentorium, excluding ventricles, with a voxel-based estimation.
    * **Left Thalamus Proper**: Represents the volume of the left thalamus, a critical relay center in the brain.
    * **BrainSegVol:** Total segmented brain volume, including both gray and white matter.
    * **SupratentorialVol:** Total supratentorial volume, including ventricles.
  This strong positive correlation suggests that raw bispectrum configurations at this scale are highly sensitive to global and regional white matter integrity and supratentorial structures.
  Intermediate angles for this configuration yield near-zero correlation across all variables. This suggests that these spatial interactions do not predominantly capture local, nor global morphological features, nor structural dependencies at mixed scales, unlike small and large angles, which emphasize self-similar structures.

Strong deviations from Gaussianity in the ventricular system (high correlation in the reduced bispectrum) suggest that ventricular expansion is associated with non-trivial morphological alterations beyond simple volumetric enlargement, possibly reflecting asymmetries, shape distortions, or interactions with surrounding structures. In contrast, a strong negative correlation between the reduced bispectrum and brain volume measures (e.g., BrainSegVol, SupratentorialVol) indicates that as brain atrophy progresses, the brain’s spatial organization deviates from a hierarchical Gaussian framework, likely due to selective tissue loss patterns.

## 6. Discussion

The findings in this work demonstrate the utility of higher-order summary statistics in understanding the relationship between brain aging and structural changes in brain anatomy and physiology. By employing two- and three-point statistics in Fourier-space, we developed a methodology capable of predicting chronological brain age while offering interpretable insights into the underlying physiological processes. This approach provides a novel avenue for uncovering age-related changes in brain structure. Furthermore, this method has the potential to enhance the accuracy of machine learning models by providing highly informative measures and effectively cleaning the data samples by detecting and removing outliers caused by measurement issues.

### 6.1. Cosmology-and-brain similarity and differences

The foundation of this work lies in an analogy between the statistical distribution of the brain and the cosmic web structure of galaxies evolving over time. Unlike cosmic surveys, where finite volume and observational limitations necessitate correction, brain volume inherently correlates with age (Fujita et al., 2023). Therefore, while skull standardization was applied to standardize volume measurements (see Appendix A), no additional corrections for the brain mask were implemented. Future work may explore more refined masking strategies.

A central insight from our analysis is the behavior of the MRI intensity power spectrum and bispectrum across different scales and ages, drawing parallels to the cosmic density field studied in cosmology. The upper left panel of Figure 3 shows the power spectrum evolution across different age groups. Similar to the growth of large-scale cosmological structures, we observe a near-linear evolution of large brain structures with increasing age.

To assess nonlinear brain evolution, we analyzed the three-point statistics. The upper right panel of Figure 3 displays the nonlinear evolution of brain structures through the reduced bispectrum for large-scale triangle configurations. Here, younger individuals exhibit greater variability in large-scale structures than older individuals, suggesting greater structural complexity in early adulthood. This aligns with previous findings on the progressive simplification of brain structure with age, partly driven by ventricular expansion. The analogy to cosmic evolution is notable: just as the universe appears more homogeneous at earlier times (Springel et al., 2005), brain structure appears to lose complexity with aging. This relationship between large-scale structural loss and brain aging supports previous studies (Yang et al., 2024; Bethlehem et al., 2022; Oschwald et al., 2019), highlighting the potential of the bispectrum as a diagnostic tool for neurodegenerative diseases.

Conversely, the raw bispectrum at small scales (bottom-right panel of Figure 3) reveals an intricate relation with age. At small and large angles, we see a decrease of structural complexity in older individuals, while at intermediate angles we see an increase of complexity with age. This is likely linked to the reduction in gray and white matter volume with age. In analogy to galaxy formation, where mergers occur in regions of high dark matter density, the reduction in brain matter may induce a “folding” or compactification of brain structures. This mirrors processes observed in cosmological phase-space evolution. As gray and white matter decrease, the corresponding brain structures may reorganize into a more intricate spatial configuration, compensating for volume loss.

### 6.2. Gender differences

Internal tests indicated that analyzing males and females separately did not improve classification sensitivity. However, using this information as an independent categorical feature in the age-regression algorithm did slightly improve the prediction. This outcome is convenient from a modeling perspective, as it avoids the need for gender-specific models. Considering that there is evidence that aging processes differ by biological gender (e.g., Liem et al., 2017), future studies with larger datasets should explore potential gender-based differences in more depth.

### 6.3. Interdisciplinary approach and future directions

The interdisciplinary framework presented here bridges cosmology and neuroscience, providing a novel perspective on brain aging. By drawing parallels to the cosmic web, where large-scale structures evolve through gravitational interactions, we gain deeper insights into how the brain undergoes structural reorganization over time. The combination of power spectrum and bispectrum methods enables us to detect features that traditional techniques may overlook.

By cross-checking the correlations between bispectra, anatomical variables, and age, we confirm key features related to brain aging, such as ventricle volume expansion, white matter reduction, and choroid plexus involvement in brain aging. These findings align with prior studies (Bethlehem et al., 2022; Dani et al., 2021; Choi et al., 2022; Danielsen et al., 2020) and reinforce that the bispectrum captures meaningful higher-order structural variations.

The bispectrum serves as a powerful tool for detecting age-related structural transformations, distinguishing between large-scale atrophy (ventricular expansion) and fine-scale changes in cortical and white matter structure. The observed scale-dependent patterns reinforce the idea that the brain undergoes a systematic transition from a more complex, interconnected state in early adulthood to a simplified, functionally reorganized state in aging.

A key advantage of the method presented here is its versatility. It can be applied beyond MRI to any imaging modality like: Positron Emission Tomography (PET) (Sweet and Brownell, 1953; Ter-Pogossian et al., 1975); Computed Tomography (CT) (Oldendorf, 1978); Single-Photon Emission Computed Tomography (SPECT) (Bruyant, 2002; Elhendy et al., 2002); Diffuse Optical Imaging/Tomography (DOI/DOT) (Martelli et al., 2009; Durduran et al., 2010) or Magnetoencephalography (MEG) (Cohen, 1968, 1972).

Future work will extend this analysis to larger datasets, incorporating functional MRI (fMRI) to explore how functional connectivity patterns evolve with age. We also aim to integrate higher-order statistics beyond the bispectrum, including the trispectrum, to capture even more nuanced structural changes. Another promising direction is longitudinal tracking of individuals to detect early markers of dementia.

Our brain age regression pipeline includes robust statistical methods for estimating uncertainties in brain age predictions. This enhances clinical applicability, making it a viable tool for medical diagnostics and research.

In analogy to cosmology, where the large-scale structure of the universe is characterized by a set of cosmological parameters, the brain might be described by a limited set of fundamental parameters that define an individual’s characteristic neuroanatomical state. Identifying such parameters could provide new biomarkers for neurological health.

This research opens avenues for improved diagnosis and treatment of neurodegenerative conditions such as Alzheimer’s, advancing our understanding of brain aging processes and providing valuable tools for clinicians and researchers alike.

## Appendix A. OASIS-3 database

Data were provided by `OASIS-3`: Longitudinal Multimodal Neuroimaging: Principal Investigators: T. Benzinger, D. Marcus, J. Morris; NIH P30 AG066444, P50 AG00561, P30 NS09857781, P01 AG026276, P01 AG003991, R01 AG043434, UL1 TR000448, R01 EB009352. AV-45 doses were provided by Avid Radiopharmaceuticals, a wholly owned subsidiary of Eli Lilly.

**Figure A.6:**
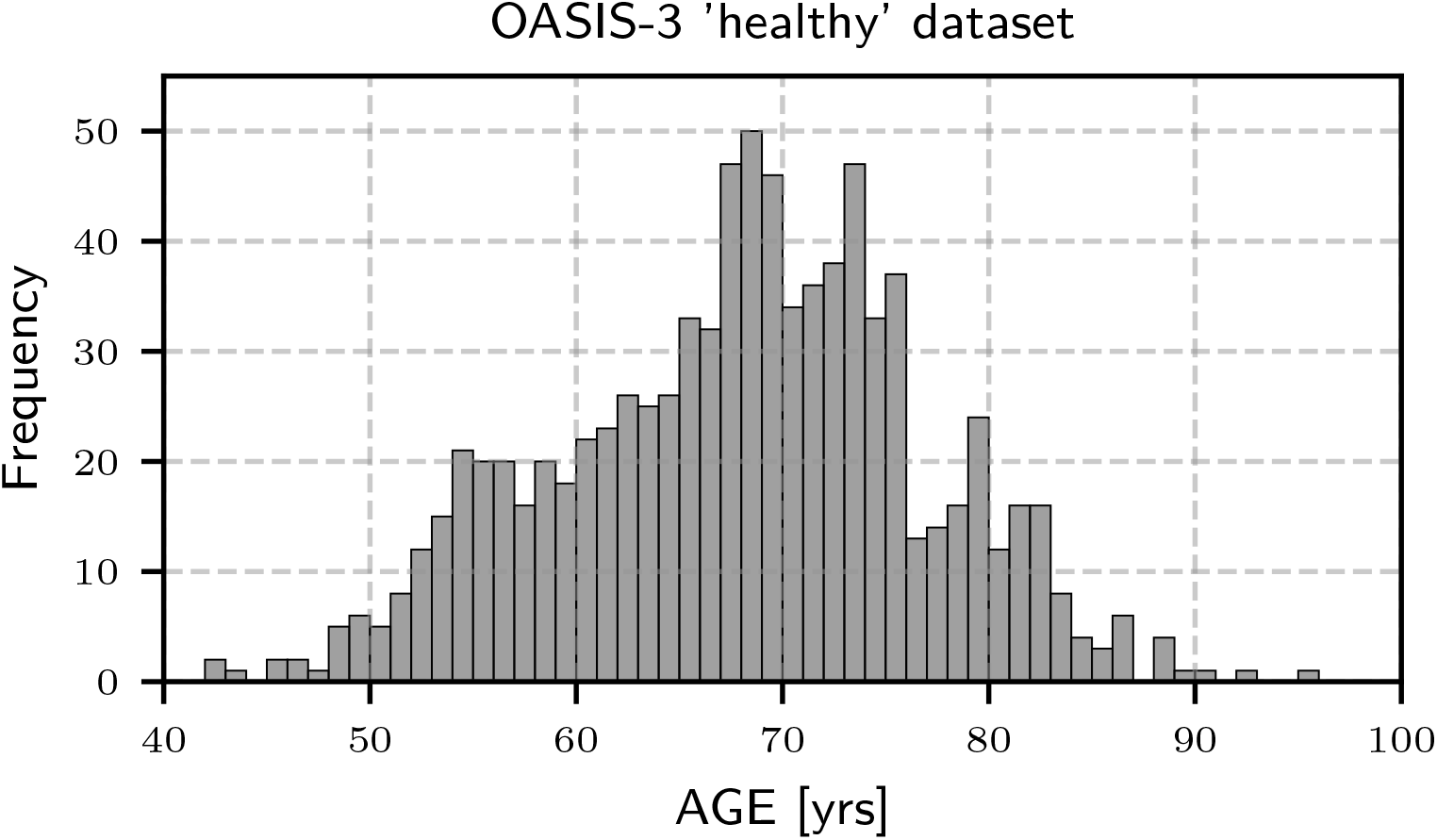
Age distribution for the OASIS-3 dataset used in the analysis.

### Appendix A.1. Selection of OASIS-3 sample

Starting from the `OASIS-3` database, we define a sample of “healthy” subjects, characterized as individuals who have not yet developed dementia (based on their Clinical Dementia Rating, CDR) and who do not suffer from any other active conditions in their clinical records. Based on this criterion, we select 401 unique participants, of whom 58% are women and 42% are men. For these “healthy” subjects, we analyze all available T1-weighted (T1w) MRI images in native space using a higher-order statistical pipeline. The dataset under study consists of 869 MRI sessions, covering a chronological age range from 42 to 98 years (see Figure A.6).

### Appendix A.2. Preprocessing of the calibration sample

We preprocess the T1-weighted (T1w) MRI data using the HCP minimal processing pipeline, as described by Glasser et al. (2013) with version `v4.3.0`. This process standardizes all images to a resolution of 1 mm, including alignment and the removal of skull and non-brain tissues. In most cases, we follow the standard HCP configuration. However, when T2-weighted (T2w) or field-map images are unavailable, we adapt the pipeline using the available imaging data, without applying the HCP-specific method to achieve reference precision.

Cortical reconstruction and volumetric segmentation were performed with the convolutional neural network `FastSurfer` suite (Henschel et al., 2020, 2022; Faber et al., 2022; Estrada et al., 2023), which performs an equivalent analysis to the widely used `FreeSurfer` (Fischl, 2012) image analysis suite. The technical details of these procedures are described in prior publications (Dale et al., 1999; Fischl et al., 1999; Fischl and Dale, 2000; Fischl et al., 2001, 2002, 2004b,a; Segonne et al., 2004; Han et al., 2006; Jovicich et al., 2006; Reuter et al., 2010, 2012).

## Appendix B. Power spectra and bispectra computations

We employ the FFTW algorithm to compute both the power spectrum and bispectrum. The process begins by transforming the dataset, originally consisting of 3D voxels in NIfTI format (with a 1mm resolution), onto a cubical mesh with side length of 300 mm and 300^3^ voxels, centered on the brain. We adapted a cosmological code (Zhao et al., 2021), traditionally used to estimate the bispectrum in cosmological simulations [in units of *h*^−1^ Mpc; 1 Mpc = 3.086 `×` 10^22^ m; *h* = *H*_0_*/*(100km*/*s*/*Mpc) with *H*_0_ being Hubble’s constant], to work with MRI data (see also Kitaura et al., 2016). This involves a direct translation from millimeters to *h*^−1^ Mpc (1 mm = 1 *h*^−1^ Mpc in numerical code units), while operating directly on MRI intensity values instead of density fluctuations as in cosmological applications. Throughout the analysis, we use a Nearest-Grid-Point (NGP) assignment kernel.

Given the intricate structure of the brain and the associated complexity across various scales, numerous configurations need to be sampled. To reduce the otherwise vast parameter space, we restrict the triangle sizes in Fourier-space to the range 0.1 *< k <* 2 mm^−1^ corresponding to spanning spatial scales from ∼ 3 to 63 mm. The lower bound excludes large-scale effects that may be dominated by boundary issues, while the upper bound, set at twice the grid resolution (1mm), helps to prevent aliasing artifacts (see, e.g., Press et al., 1992; Jing, 2005).

## Appendix C. Metrics used in this study

- **Bispectrum variance**: 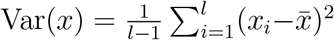 with 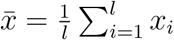, where *x*_*i*_ is the bispectrum value for an angle *θ* at bin *i* and *l* being the total number of *θ* bins.
- **MAE (mean absolute error)**: 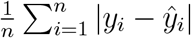 with *y*_*i*_ and *ŷ*_*i*_ being the predicted and true age values for session *i* out of *n* sessions, respectively. The data array vector conformed by all predicted ages is given by *Y*, while for the true ages it is *Ŷ*.
- **Geometric Mean (G-Mean)** for the classification performance is calculated as the geometric mean of the sensitivities across all age groups: 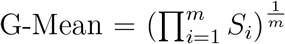, where *m* is the total number of age groups or classes, and: 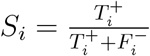 is the sensitivity (recall) for class (age group) *i, T*^+^ are the true positives and *F* ^−^ are the false negatives. A true positive occurs when the model correctly predicts that a subject belongs to their true age group. A false negative occurs when a subject truly belongs to a specific age group, but the model predicts them to be outside that group.
- **Kendall’s correlation coefficient** calculation steps:
  1. Rank the subjects: Rank all subjects based on their bispectrum values for the specific configuration we are analyzing. Separately, rank the same subjects by their biological ages.
  2. Compare pairs of subjects: For every pair of subjects, determine if the ranks of both variables (bispectrum and biological age) are consistent:
    - Concordant pair: If one subject has a higher bispectrum value and is also older, or if one has a lower bispectrum value and is also younger.
    - Discordant pair: If one subject has a higher bispectrum value but is younger, or has a lower bispectrum value but is older.
  3. Compute 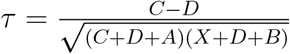, with *C* and *D* being the number of concordant and discordant pairs, respectively; *A* the number of ties only in *Y*, and *B* the number of ties only in *Ŷ*.
- Covariance matrix: Let 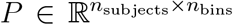 be the matrix of age distributions for *n*_subjects_ number of subjects and *n*_bins_ number of bins (we consider a binning of 4 years), where each row *p*_*s*_ is a probability distribution over bins for subject *s*. Then the (unnormalized) covariance matrix is:

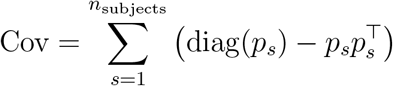

The normalized covariance matrix is computed as:

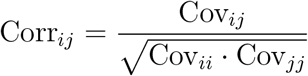

with the convention that Corr_*ij*_ = 0 if the denominator is zero.

## Appendix D. Summary of the method

The following steps are outlined in Figure 2:

1. **MRI data preparation:** We start with the selection and preprocessing of MRI data and representation of intensity distribution on a cubical regular mesh (see Appendix A).
2. **Higher-order statistical three-dimensional spatial frequency analysis**
  a. **Fourier-space calculations:** We Fourier transform each MRI data cube and compute its power spectrum and bispectra for a large variety of configurations. The power spectra are used to construct the hierarchical models, which in turn are used to compute the reduced bispectra (see bottom panel in Figure 2 and Appendix B).
  b. **Pre-classification scale-dependent analysis of systematics:** We analyze the variance of the bispectra across different age groups. Outliers, likely due to dementia or MRI measurement issues, exhibit significant deviations from the average variance within their respective age group. To ensure the integrity of the dataset, we apply a conservative exclusion criterion. Initially, we focus on the raw and reduced bispectrum configurations with the smallest variance variations across age, in order to better address systematics. The configurations identified are (*k*_1_ = 0.13, *k*_2_ = 0.57 mm^−1^) for the raw bispectrum and (*k*_1_ = 0.59, *k*_2_ = 0.80 mm^−1^) for the reduced bispectrum. Next, we exclude sessions that deviate by more than ≳3 standard deviations from the mean variance. As a result, we reduce the dataset from 921 sessions to 880 using the reduced bispectrum and to 869 using the raw bispectrum.
  c. **Classification and scale-dependent correlation analysis:** We analyze Kendall’s rank correlation coefficients between configurations and age (see Figure D.7). Additionally, we apply a Random Forest classifier, computing the G-mean and MAE for each singular bispectrum configuration. The configurations with the highest G-means and lowest MAEs are identified and considered. These top-performing configurations are then combined in all possible ways to reduce the selection to a minimal set, capturing the most important characteristic scales. The Kendall coefficients support our choice of pre-classification analysis for systematics. According to *f* (*k*_1_, *k*_2_), the raw bispectrum with the lowest correlation with age is ≲ 40000 for *k*_1_ ≲ 0.4 mm^−1^, while for the reduced bispectrum, it is ≳ 50000 for *k*_1_ ≳ 0.5 mm^−1^.
  d. **Neuro-anatomical and physiological interpretation:** The proposed method enables physiological interpretation by examining correlations between the summary statistics, age, and anatomical variables. We employed Kendall’s rank correlation to capture potential nonlinear associations. For anatomical features, we used regions defined by `FreeSurfer`. Our analysis reproduced well-established aging effects, including grey and white matter atrophy and ventricular enlargement.
3. **Age regression**
  a. **Neural network design:** We develop an age-regression neural network integrating the selected bispectrum configurations and gender into a multi-input architecture. Each configuration is processed through dense layers employing ReLU activations. The resulting features are concatenated and passed through fully connected layers with batch normalization, regularization and dropout to improve generalization. The final output layer also applies a ReLU activation. The model is trained using huber loss and optimized with Adam schedule.
  b. **Deep ensemble:** In each realization we train and predict the dataset using a stratified k-folding method. This way, we predict the age for all MRI sessions in the dataset. Finally, we run 1000 realizations to estimate a deep ensemble from where we estimate the mean as the predictor and the standard deviation as the error.

**Figure D.7:**
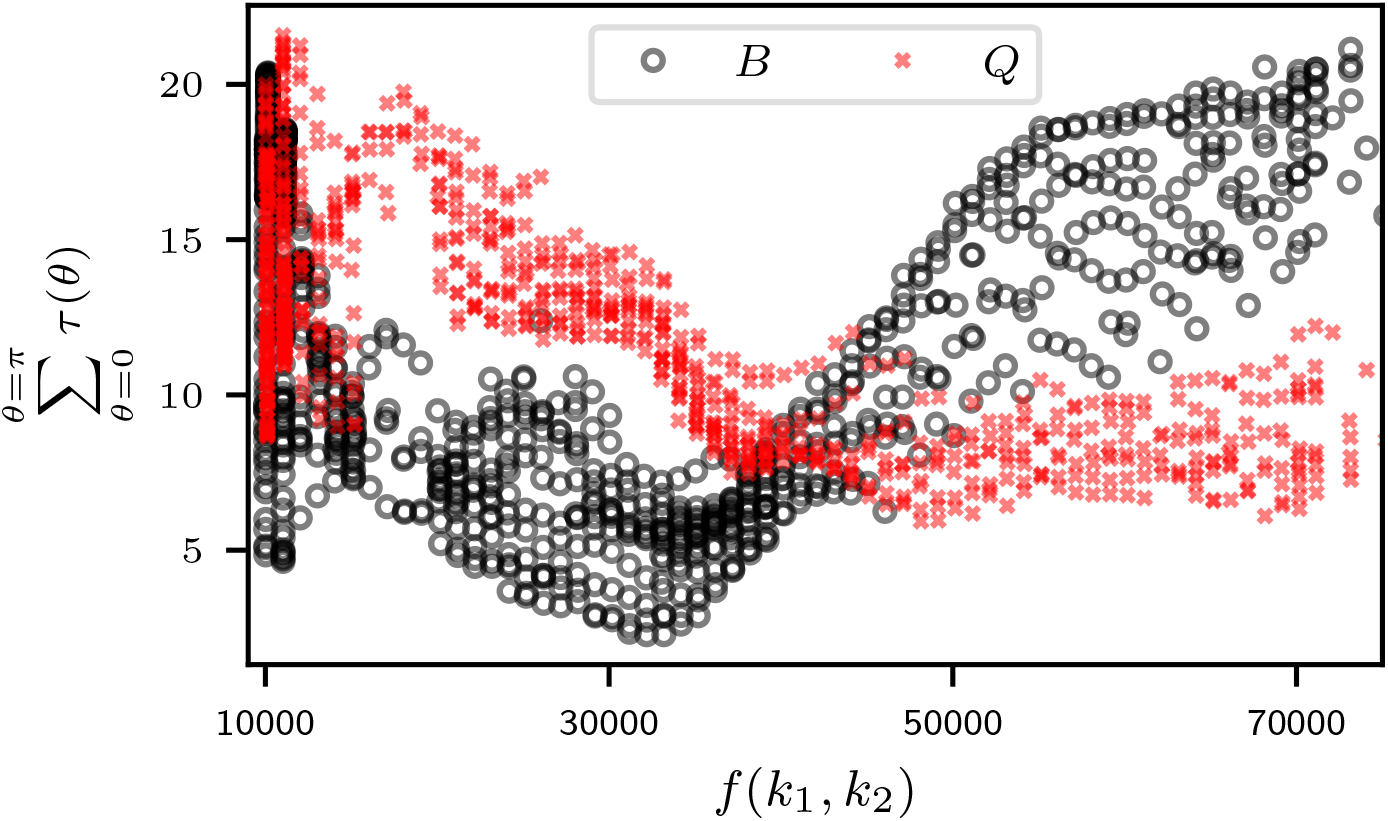
Sum of Kendall’s rank correlation coefficient between configurations and age for the `OASIS-3` sample. The *x*-axis represents a combination of *f* (*k*_1_, *k*_2_) = 100000 × *k*_1_ + 100 × *k*_2_, while the *y*-axis shows the sum of *τ* within *θ*-bins of 2^*°*^ 15^*′*^ arcminutes. The trend at ∼ 70000 continues to higher *k*-configurations. This representation highlights the dependence on *k*_1_: *k*_1_ ∼ *f* (*k*_1_, *k*_2_)*/*10^5^, e.g., *f* (*k*_1_, *k*_2_) = 50000 : *k*_1_ ∼ 0.5.

## Appendix E. Validation on MRI simulations

Following Pinaya et al. (2022), we applied a Latent Diffusion Model – downloaded as a pre-trained model from the Medical Open Network for Artificial Intelligence project (MONAI Consortium, 2021; Cardoso et al., 2022)– conditioned on age, gender, ventricle size, and brain volume to generate synthetic MRI images. This approach yielded 2,235 images with characteristics closely resembling those of the `OASIS-3` dataset.

**Figure E.8:**
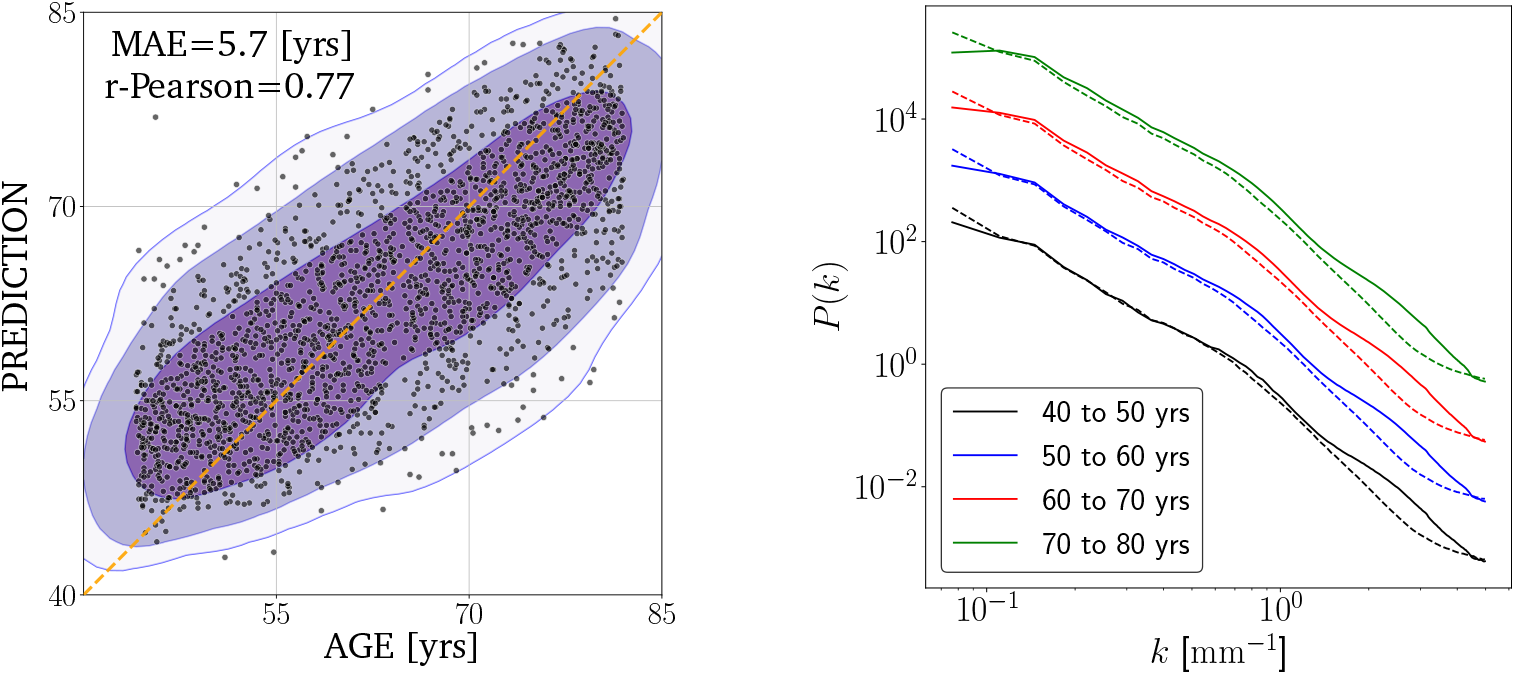
Left panel: age regression for the simulated `OASIS-3` dataset. Using the deep-ensemble technique, we estimate the age for all MRI sessions, resulting in a MAE of 5.5 years. Right panel: power spectra comparison for real (solid line) vs simulated data (dashed line). For visual purposes we apply a multiplicative factor to each age power spectra of 1,10,100 and 1000 respectively.

Images were skull stripped following the `pre-Freesurfer` step from the HCP preprocessing pipeline and then run through the age-inference method. The results are shown in Figure E.8. The model’s performance on the synthetic data was substantially inferior to its performance on real data.

To investigate the cause of this performance gap, we computed the power spectra across different age ranges, comparing the simulations to real data. We observed good agreement in the low-frequency range (*k <* 1), but a marked discrepancy at higher frequencies. The simulated data exhibited a lack of power at small scales, suggesting that fine-grained features are not accurately captured. Additionally, the large-scale power diverged increasingly with age, indicating that age-related structural changes are not completely well represented in the simulations. Our results demonstrate that higher-order statistical analysis provides a valuable tool for assessing the accuracy of simulations, analogous to methods employed in cosmological studies (e.g., Springel et al., 2005; Kitaura et al., 2016).

## Appendix F. Application to the Cam-CAN dataset

We consider the `Cam-CAN` dataset. MRI data for `Cam-CAN` participants were acquired on a 3 T Siemens TIM Trio scanner equipped with a 32-channel head coil. T1-weighted anatomical images were collected using a 3D MPRAGE sequence (TI/TR/TE = 900/2250/2.99 ms, flip angle = 9°, FOV = 256 × 240 × 192 mm, 1 mm isotropic voxels, GRAPPA = 2; acquisition time = 4 min 32 s).

**Figure F.9:**
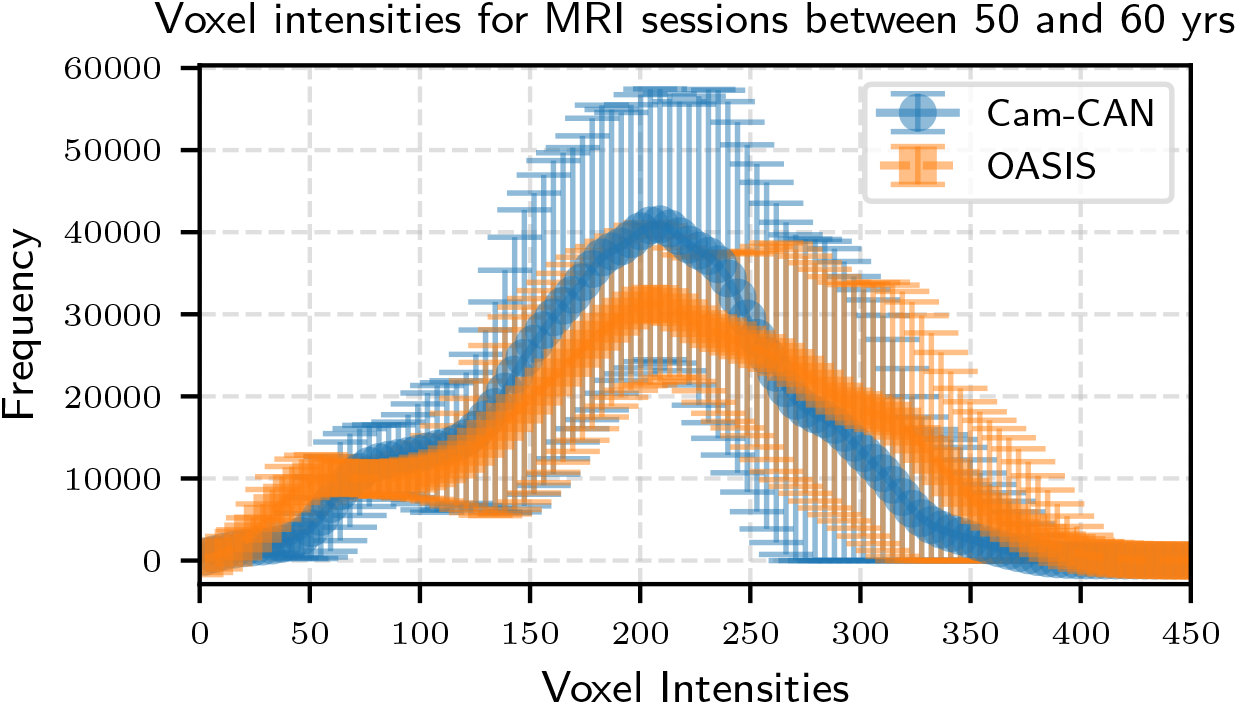
Voxel intensity distributions and corresponding standard deviations for MRI sessions aged between 50 and 60 years from the `OASIS-3` and `Cam-CAN` datasets. The differences observed within this age range are consistent across all other age groups.

Systematic differences between the `OASIS-3` and `Cam-CAN` datasets, clearly illustrated by intensity distribution inconsistencies shown in Figure F.9, prevented us from applying the previously identified `OASIS-3` bispectrum configurations directly to the `Cam-CAN` dataset. Consequently, we conducted an analogous configuration selection study specifically for `Cam-CAN`. The optimal configurations identified differ in two configurations from those derived for `OASIS-3`, most notably, the model favors the use of the power spectrum instead of the reduced bispectrum. In Table F.2 we summarize the configurations used. Median values as a function of age, for `Cam-CAN` configurations are found in Figure F.10.

We processed the dataset with the same HCP Pipeline and remove outliers based on discrepant variances on configurations `B080116` and `B010045`, resulting in 625 MRI sessions, spanning an age range from 18 to 88 years (shown in Figure F.11) with a much lower density of subjects per age. Next, we repeated the search for the best configurations, ending with the list shown in Table F.2.

**Figure F.10:**
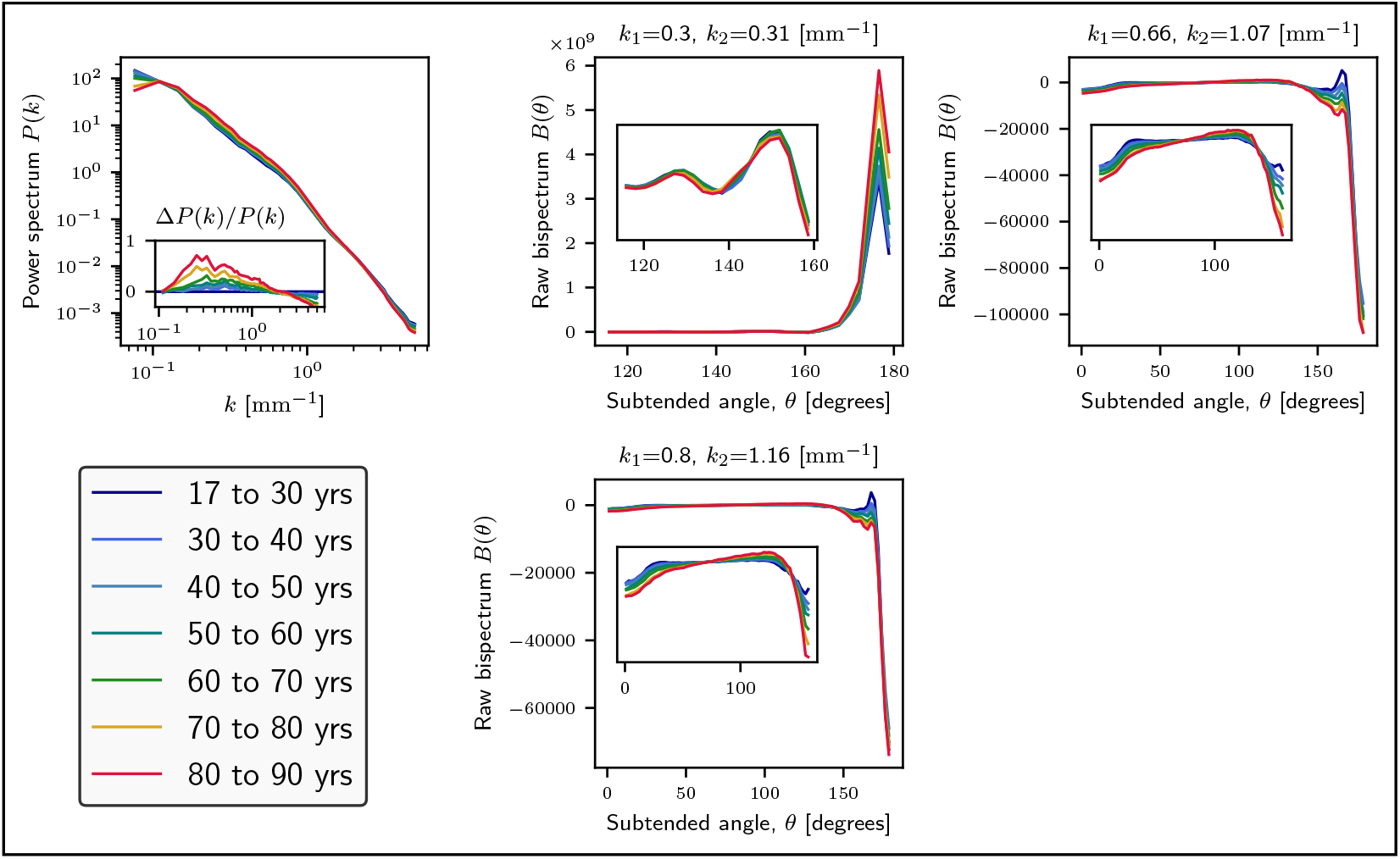
Median power spectra and bispectra per age group for the individual configurations that enter the biomarker definition from the `Cam-CAN` dataset. Here we consider the raw bispectra for fix |**k**_1_| and |**k**_2_| as a function of the subtended angle *θ* between both *k*-vectors: *B*(*θ*|*k*_1_, *k*_2_).

**Figure F.11:**
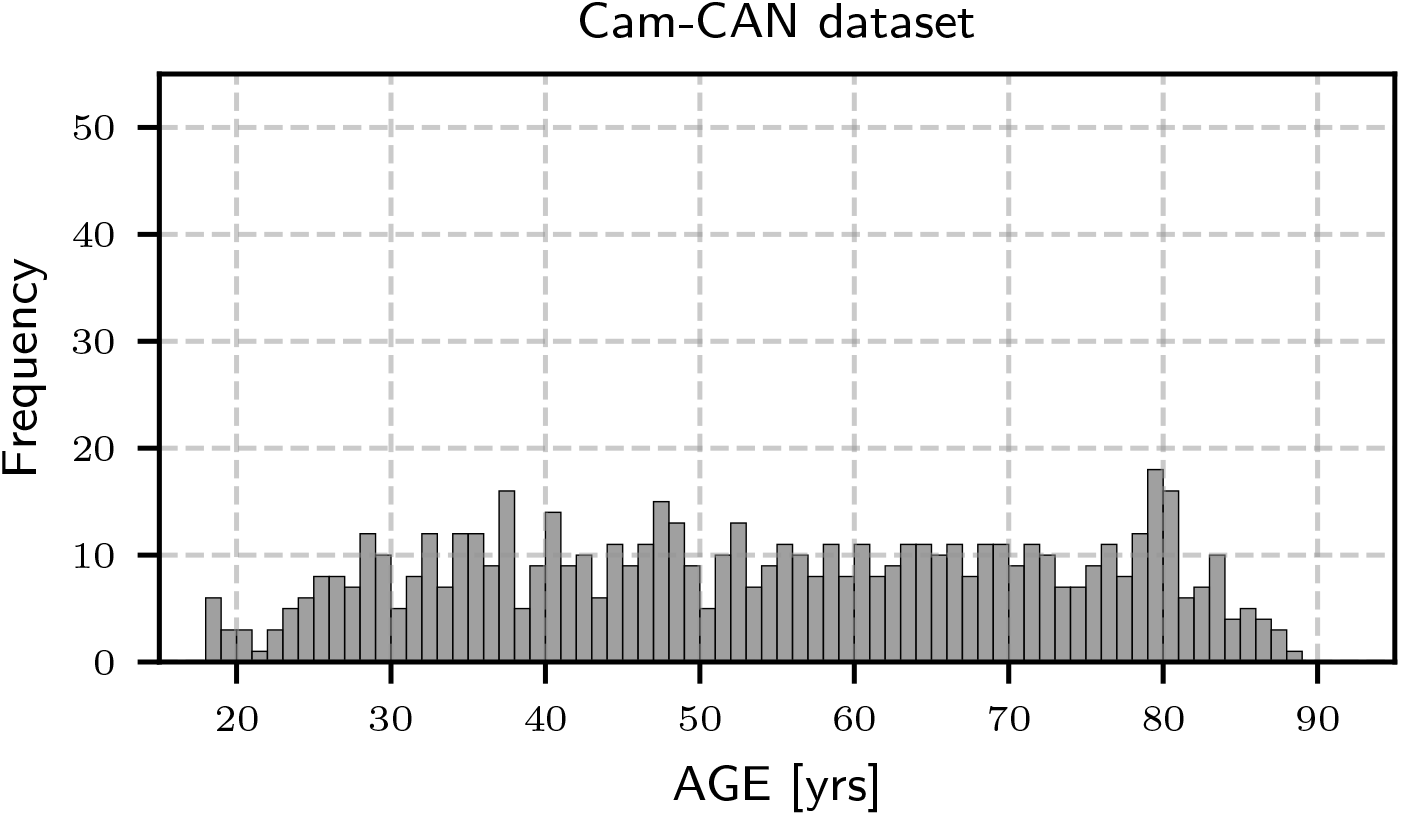
Age distribution for the `Cam-CAN` dataset. The frequency axis is in the same scale as in Figure A.6, highlighting the density differences with respect to `OASIS-3` dataset.

**Table F.2:** Summary of singular configurations making the `Cam-CAN` biomarker

| | $k_1$ [mm<sup>-1</sup>] | $k_2$ [mm<sup>-1</sup>] | $r_1$ [mm] | $r_2$ [mm] | $\theta$ [rad] | Name |
| --- | --- | --- | --- | --- | --- | --- |
| $B$ | 0.30 | 0.31 | 20.94 | 20.27 | 2.-3.15 | B030031 |
| $B$ | 0.66 | 1.07 | 9.52 | 5.87 | 0.-3.15 | B066107 |
| $B$ | 0.80 | 1.16 | 7.85 | 5.43 | 0.-3.15 | B080116 |
| $P(k)$ | | | | | | P(k) |

At last, we applied the age-regression neural network to the `Cam-CAN` dataset, achieving a MAE of approximately 5.9 years (see Figure F.12). The MAE we obtain is moderately higher than that achieved by machine learning methods directly applied to structural MRI data, such as in Xifra-Porxas et al. (2021), who report a MAE of 5.33 years after optimization. Whether this discrepancy arises from statistical fluctuations or reflects the limitations of using two-point and three-point statistics in the considered scales to represent brain structure remains an open question that warrants further investigation. Incorporating four-point statistics, or accounting for anisotropic clustering—e.g., through a Legendre multipole expansion of the correlation function—may help capture additional, relevant information.

The right panel of Figure F.12 highlights the compatibility between the `OASIS-3` and `Cam-CAN` results, suggesting the potential for a joint analysis across datasets. While appealing, this is far from straightforward: not only do the probability density functions (PDFs) of the two datasets differ substantially (see Figure F.9), but their respective power spectra also show significant discrepancies, as evident from the power spectrum ratio plots in Figures 3 and F.10. These differences indicate that some form of preprocessing or harmonization is required to enable the use of shared bispectrum configurations. As an alternative, a joint analysis could be pursued by explicitly categorizing or stratifying the datasets, allowing the model to account for systematic differences between them. We leave these studies for future work.

**Figure F.12:**
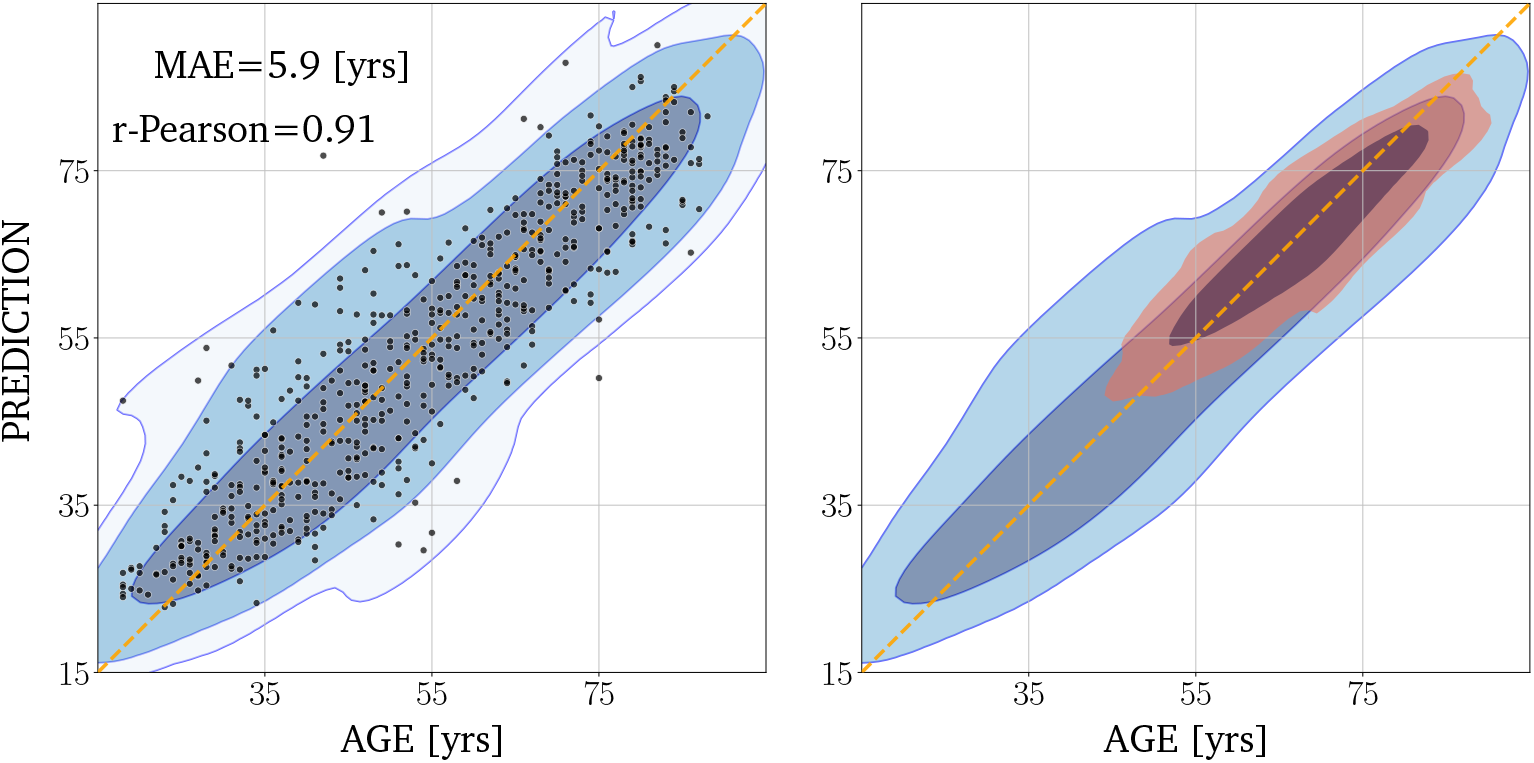
Age regression for the `Cam-CAN` dataset. On the left: using the deepensemble technique, we estimate the age for all MRI sessions, resulting in a MAE of 5.9 years. On the right: one- and two-*σ* contour regions for the `OASIS-3` (in red) and the `Cam-CAN` (in blue) datasets.

## Acknowledgements

This research was primarily funded by the Cabildo de Tenerife through the IACTEC Technological Training Program, under the TF INNOVA grant. The funding was provided within the framework of MEDI-FDCAN 2016–2025, as part of the Tenerife Island Strategic Development Plan, in support of the `Cosmic Brain` project initiated in 2021 (Principal Investigator, PI: FSK). The authors acknowledge the Spanish Ministry of Science and Innovation for financing the `Big Data of the Cosmic Web` project: PID2020-120612GB-I00/AEI/10.13039/501100011033 (PI: FSK) and the `Cosmology with Large Scale Structure Probes` project funded by the IAC (PI: FSK), which provided numerical tools and computing facilities. NJ thanks financial support from PID2021-127611NB-I00 (PI: NJ).

The authors also wish to acknowledge the contribution of the IAC High-Performance Computing support team and hardware facilities to the results of this research.

## Ethics statement

This study used publicly available data from the `OASIS-3` and `Cam-CAN` datasets. `OASIS-3` (LaMontagne et al., 2019) is a de-identified, open-source compilation of neuroimaging and clinical data collected through the Washington University Knight ADRC. All procedures involving human participants in `OASIS-3` were approved by the Institutional Review Boards of Washington University in St. Louis, and written informed consent was obtained from all participants. All analyses were conducted in compliance with the `OASIS-3` Data Use Agreement, which prohibits attempts to re-identify participants and requires acknowledgment of the data source in all publications.

`Cam-CAN` (Taylor et al., 2017; Shafto et al., 2014) is a population-based, multimodal (MRI, MEG, and cognitive-behavioural) dataset collected by the MRC Cognition and Brain Sciences Unit and University of Cambridge, comprising a cross-sectional adult lifespan sample (ages 18–87). The data were collected in compliance with the Declaration of Helsinki and approved by the Cambridgeshire 2 Research Ethics Committee (reference: 10/H0308/50). Written informed consent was obtained from all participants.

As both datasets are publicly available and fully de-identified, the present secondary analyses did not require additional institutional ethics approval. No additional data collection involving human participants was performed by the authors.

The study was approved by the Research Ethics and Animal Welfare Committee (CEIBA) at the University of La Laguna (CEIBA2017-0270) obtained by NJ.

## Data and code availability statement

### Data

This study uses publicly available MRI data from two cohorts: (i) `OASIS-3` (Open Access Series of Imaging Studies) and (ii) `Cam-CAN` (Cambridge Centre for Ageing and Neuroscience). Access to both datasets is available to qualified researchers upon registration and agreement to the respective data-use terms. Because of these agreements and participant privacy, we cannot redistribute raw images. No new raw MRI data were collected.

### Code

A GitHub repository contains all bispectrum calculations used in the brain age regression, together with the corresponding neural network code: https://github.com/aureliocarnero/brainage-higherorder/.

## Declaration of generative AI and AI-assisted technologies in the writing process

During the preparation of this work the author(s) used Writefull and ChatGPT to enhance the clarity and readability of the manuscript. After using this tool/service, the author(s) reviewed and edited the content as needed and take(s) full responsibility for the content of the publication.

## Footnotes

1 www.cosmic-brain.org

